# NRP1-expressing monocyte/macrophages promote arteriogenesis in the ischemic limb

**DOI:** 10.1101/2024.02.11.579823

**Authors:** JS Cho, AS Patel, A Fantin, FE Ludwinski, CJ Pichette, M. Rudnicki, L Denti, A Mondragon, P Saha, A Smith, C Ruhrberg, B Modarai

**Author notes:** **Co-corresponding authors:** Professor Bijan Modarai, Academic Department of Vascular Surgery, St Thomas’ Hospital, School of Cardiovascular Medicine and Sciences, 1st Floor North Wing, London SE1 7EH, Professor Christiana Ruhrberg, UCL Institute of Ophthalmology, 11-43 Bath Street, London EC1V 9EL. joint first author (JS Cho, AS Patel).

## Abstract

Peripheral arterial disease can cause limb threatening blood flow restriction, requiring amputation in up to a third of patients despite contemporary treatments. Stimulating new blood vessel growth in ischemic limbs using growth factors such as VEGF-A or supplying mixed cell populations via autologous transplantation has been investigated as a therapy for surgically intractable cases, but to date has proven ineffective in randomised controlled clinical studies. To identify more potent pro-arteriogenic cell populations, we have investigated VEGF-A responsive myeloid cell populations in a mouse model of severe hindlimb ischemia. We found that NRP1 ablation from monocytes and macrophages impaired arteriogenesis in the adductor muscle and flow recovery in the injured limb. Vice versa, the exogenous delivery of NRP1-expressing (but not NRP1-negative) macrophages enhanced arteriogenesis in the adductor and flow recovery in the ischemic limb in a VEGF signalling-dependent manner. Further, patient-derived NRP1-expressing monocytes promoted vascular morphogenesis ex vivo. As the levels of circulating NRP1-expressing monocytes were raised in patients with limb ischemia, the therapeutic delivery of autologous NRP1-expressing monocytes could be explored as a treatment for critical limb ischemia.

## Introduction

Therapeutic neovascularization aims to stimulate new vessel growth in ischemic tissues by angiogenesis and arteriogenesis and has been investigated as a therapy for intractable limb or myocardial ischemia. All neovascularization strategies attempted thus far, however, have failed to demonstrate efficacy in randomised controlled studies; accordingly, critical limb ischemia (CLI), the most severe manifestation of peripheral arterial disease, still results in amputation in almost a third of patients one year after diagnosis^1–4^. Investigated therapeutic strategies have included intramuscular delivery of proangiogenic factors, such as the vascular endothelial growth factor VEGF-A^5^ or fibroblast growth factor FGF2^6^, and injections of cells thought to promote neovascularization. Most studies to date have delivered a mixed population of autologous cells that include mononuclear cells derived from either bone marrow or peripheral blood, endothelial progenitor cells, hematopoietic stem cells, bone marrow concentrates, or mesenchymal stromal cells^7–12^. However, the precise effects of the different cell types after transplantation remain unclear, and it is conceivable that only some of the delivered cell types will have the desired pro-angiogenic/arteriogenic properties, whilst other cells may instead be ineffective or even prevent angiogenesis and arteriogenesis^13^. Accordingly, it is now acknowledged that mixed populations of autologous cells have insufficient angiogenic and/or arteriogenic properties for a therapeutic effect and should be substituted by subpopulations of cells with angiogenic and/or arteriogenic potency that are more likely to promote effective neovascularization when delivered to ischemic tissues in large numbers^14^. As the intramuscular delivery of such cells has the potential to benefit not only patients with critical limb ischemia but also treat those with myocardial ischemia, it has become important to identify such pro-angiogenic subpopulations with angiogenic and/or arteriogenic potency.

Monocytes and macrophages promote neovascularization of ischemic tissues in preclinical studies but typically also comprise heterogeneous cell populations^15–18^. Identifying potent pro-angiogenic/arteriogenic myeloid subsets may therefore allow the development of a clinically efficacious therapeutic product. Moreover, defining their mechanism of action may allow the identification of biologics that promote the selective recruitment of these beneficial myeloid cell types or mimic their mode of action. Further, identifying how ischemia-driven VEGF-A expression functionally intersects with monocyte and macrophage roles should help design effective therapies that promote neovascularisation. For example, prior work showed that transgenic VEGF-A induction recruits circulating monocytes to the ischemic limb and facilitates their education into pro-arteriogenic monocytes^19^. Yet, the monocyte VEGF-A receptor FLT1 (VEGFR1) is dispensable for this pathway^19^. Further work is therefore required to establish whether VEGF-A can signal directly to monocytes or macrophages to promote ischemic neovascularisation, and whether VEGF-A responsive monocytes or macrophages could be exploited therapeutically to treat limb ischemia.

NRP1 is a transmembrane glycoprotein that binds to VEGF-A as well as class 3 semaphorin (SEMA3) proteins^20–22^. In particular, an alternatively spliced isoform of VEGF-A termed VEGF165 promotes complex formation of NRP1 and VEGFR2 (KDR)^23^ to enhance MAP kinase signalling in endothelial cell cultures via ERK1/2 (MAPK3/1) and p38 (MAPK14)^24^ ^25^. Studies with genetically modified mice lacking NRP1’s cytoplasmic domain (NCD) showed that pro-arteriogenic, VEGF165-induced ERK signalling depends on the NCD-binding GIPC1 adaptor protein (also known as synectin or NRP1 interacting protein, NIP)^26^. Specifically, GIPC1 tethers the VEGF165-activated VEGFR2/NRP1 complex to myosin VI to promote its endocytosis, whereby endosome trafficking prolongs VEGFR2 signalling via ERK1/2 by increasing its protection from dephosphorylation by PTP1B^26–28^. Accordingly, the NCD is required for arteriogenesis and blood flow recovery in a mouse model of hindlimb ischemia (HLI)^26^. Although the NCD requirement for HLI recovery has been attributed to NRP1 roles in endothelial cells, the injection of NRP1-expressing monocytes and macrophages into growing tumours also promotes arteriogenesis^29^. Moreover, adenoviral overexpression of VEGF-A induces the recruitment of myeloid cells to non-ischemic muscle in a NRP1-dependent mechanism^30^. Yet, it is not known whether NRP1-expressing monocytes or macrophages promote angiogenesis or arteriogenesis in ischemic muscle, and whether such a role would involve the NCD or MAPK signalling.

Here, we have combined gene expression, cell therapy and genetic knockout studies with a mouse model of severe HLI to investigate whether NRP1-expressing monocytes or macrophages promote revascularization of ischemic muscle and whether their intramuscular delivery could enhance arteriogenesis in an NCD-dependent mechanism. Further, we have examined whether circulating NRP1-expressing monocytes isolated from patients with limb ischemia have the potential to stimulate angiogenesis and arteriogenesis *in vitro*.

## Results

### NRP1-expressing monocytes and macrophages increase after hindlimb ischemia

Unilateral ligation and excision of the femoral artery has previously been used as a mouse model to evaluate recovery from critical limb ischemia^31^. Analysis of a publicly available transcriptomic dataset from ischemic adductor muscle obtained using this model^32^ demonstrated upregulation of the monocyte/macrophage markers *Cd68*, *Csf1r* (CD115) and *Adgre1* (F4/80) in the ischemic adductor muscle between day (D) 1 and D14 after surgery (**Fig. 1A**). On D1, we also observed upregulation of *Il6*, a marker for M1-like, pro-inflammatory macrophages, which was followed on D7 after surgery by upregulation of *Mrc1*, a marker for M2-like, tissue repair-promoting macrophages, the latter coinciding with upregulation of the pan-endothelial markers *Pecam1* and *Cdh5* and the arterial marker *Gja4* (**Fig. 1B**). Moreover, the dual endothelial and monocyte/macrophage marker *Nrp1*^29,30,33^ was upregulated from D1 after surgery onwards (**Fig. 1A**). Together, these finding correlate the presence of M2 macrophage markers and NRP1 expressing with the onset of arteriogenesis.

**Fig. 1.**
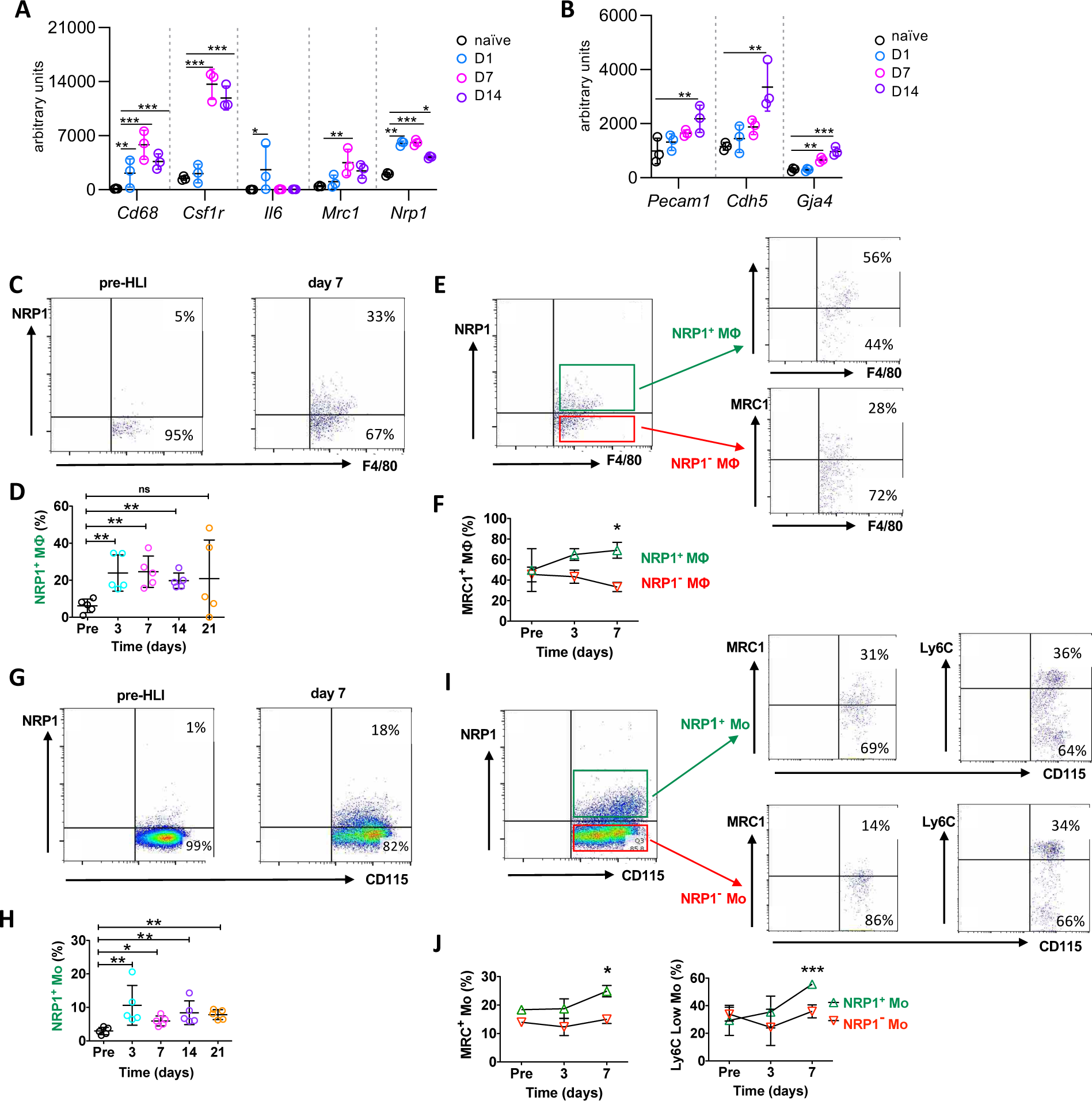
NRP1-expressing monocytes and macrophages increase after hindlimb ischemia. (**A,B**) *Increased expression of myeloid and vascular markers in ischemic muscle*. Analysis of publicly available microarray data for the indicated myeloid markers and *Nrp1* (**A**) and the indicated endothelial markers (**B**) in ischemic wild-type hindlimb adductor muscle at the indicated time-points relative to naïve muscle; n = 3 mice/group/time point; data are shown as mean ± SD; 2-way ANOVA (P = <0.0001) and Dunnett’s multiple comparison test. (**C-F**) *NRP1-expressing macrophages with a tissue-remodelling phenotype increase in the ischemic adductor muscle*. Macrophages were identified by flow cytometry as CD45^+^ CD11b^+^ F4/80^+^ Lin^-^ cells in the ischemic muscle of wild-type mice at the indicated time points before and after surgical HLI induction. (**C,D**) Proportion of NRP1^+^ macrophages amongst all macrophages, including example dot plot analyses (**C**) and quantification (**D**); each data point represents the value for one mouse; n = 5 mice/group/time-point; data are shown mean ± SD; Mann-Whitney test between time-points. (**E,F**) Proportion of MRC1^+^ macrophages amongst NRP1^+^ and NRP1^-^ macrophages, including example dot plot analyses (**E**) and quantification (**F**); day 7: MRC1^+^ NRP1^+^ 69.1 ± 7.8% vs. MRC1^+^ NRP1^-^ 33.3 ± 4.5%; n = 5 mice/group/time-point; data are shown mean ± SEM; 2-way ANOVA P = 0.02, followed by Bonferroni multiple comparison test. (**G-I**) *NRP1-expressing monocytes with a tissue-remodelling phenotype increase in the circulation after HLI induction.* Monocytes were identified by flow cytometry as CD115^+^ CD11b^+^ Lin^-^ cells in peripheral blood of wild-type mice at the indicated time points before and after surgical HLI induction. (**G,H**) Proportion NRP1^+^ monocytes amongst all monocytes, including example dot plot analyses (**G**) and quantification (**H**); each data point represents the value for one mouse; n = 5 mice/group/time-point; data are shown mean ± SD; Mann-Whitney test to compare time-points. (**I,J**) Proportion of MRC1^+^ and Ly6^low^ monocytes amongst NRP1^+^ and NRP1^-^ monocytes, including example dot plot analyses (**I**) and quantification (**J**); each data point represents the mean ± SEM of n = 5 mice/group/time point. Day 7: MRC1^+^ NRP1^+^ 24.9 ± 2.0% vs. MRC1^+^ NRP1^-^ 15.0 ± 1.5%; Ly6C^low^ NRP1^+^ 55.6 ± 1.2% vs. Ly6C^low^ NRP1^-^ 35.9 ± 2.1%; 2-way ANOVA P = 0.004 (MRC1) and P = 0.003 (Ly6^low^), followed by Bonferroni multiple comparison test. Abbreviations: MΦ, macrophage; Mo, monocyte; ns, not significant (P >0.05), * P ≤0.05, ** P ≤0.01, *** P ≤0.001.

Flow cytometric analysis of the adductor following HLI surgery in wild type mice confirmed that increased NRP1 levels could be, at least in part, explained by an increase in NRP1-expressing macrophages, because their proportion from D3 up to D14 after injury increased significantly in ischemic muscle when compared to NRP1-negative macrophages (**Fig. 1C,D**; Supplementary **Fig. S1A**). Also agreeing with the expression analysis, a greater proportion of the muscle-resident NRP1-expressing macrophages expressed MRC1 on D7 after surgery when compared with NRP1-negative macrophages (**Fig. 1E,F**). NRP1 expressing macrophages with M2-like characteristics are therefore enriched in ischemic muscle at the onset of arteriogenesis.

Flow cytometric analysis further showed that the proportion of NRP1-expressing monocytes in peripheral blood increased significantly from D3 to at least D21 after injury (**Fig. 1G,H**; **Supplementary Fig. S1B**). There was a greater proportion of Ly6C^low^ and MRC1-positive cells in the D7 population of NRP1-expressing monocytes compared with NRP1-negative monocytes (**Fig. 1I,J**). These findings agree with prior work demonstrating that MRC1-expressing monocytes are upregulated in hindlimb ischemia^34^, and that pro-angio/arteriogenic, MRC1-expressing and Ly6C^low^ monocytes, respectively, accumulate during tumour angiogenesis^35^ or after myocardial infarction^36^. Taken together, marker analysis by transcriptomic and flow cytometry analysis showed that femoral artery ligation increased the proportion of NRP1-expressing monocytes and macrophages in the circulation and ischemic muscle of the mouse.

### NRP1-expressing macrophages promote revascularization of ischemic limbs

We next asked whether NRP1-expressing macrophages promote recovery from ischemia in the HLI mouse model. For a loss of function experiment, we bred mice carrying Cre recombinase expressed from the endogenous myeloid promoter for *Lyz2*, also known as *Lysm*, to mice carrying *Nrp1* floxed allele, thus generating mice that lack NRP1 in monocytes and macrophages (*Lysm^Cre/+^;Nrp1^fl/fl^*)^37^ and control mice without the Cre knock-in allele (*Lysm^+/+^;Nrp1^fl/fl^*, referred to as controls). In agreement with prior literature^37^, flow cytometry of naïve mice that also carried *Lysm^Cre/+^* alongside the *Rosa^Yfp^* recombination reporter showed that on average ∼55% of circulating monocytes were targeted with YFP and, accordingly, NRP1-expressing monocytes in *Lysm^Cre/+^;Nrp1^fl/fl^* mutant mice appeared reduced by ∼50% (**Fig. 2A**). The reduced percentage and number of NRP1-expressing macrophages was even more evident (∼90% reduction) in bone marrow-derived macrophages (BMDM) from *Lysm^Cre/+^;Nrp1^fl/fl^*mutants compared to controls (**Fig. 2B**). Doppler analysis showed that *Lysm^Cre/+^;Nrp1^fl/fl^* mutants had significantly impaired flow recovery in the ischemic limb following HLI compared with controls, in particular, from D7 after surgery onwards (**Fig. 2C,D**). Quantitative analysis of immunostained sections from ischemic adductor muscle on D28 after surgery showed significantly reduced arteriole density and arteriole diameter in *Lysm^Cre/+^;Nrp1^fl/fl^*mice compared with controls (**Fig. 2E,F**). Together, these findings suggest that NRP1-expressing macrophages are required for arteriogenesis in ischemic muscle tissue and therefore for flow recovery after induction of ischemia.

**Fig. 2.**
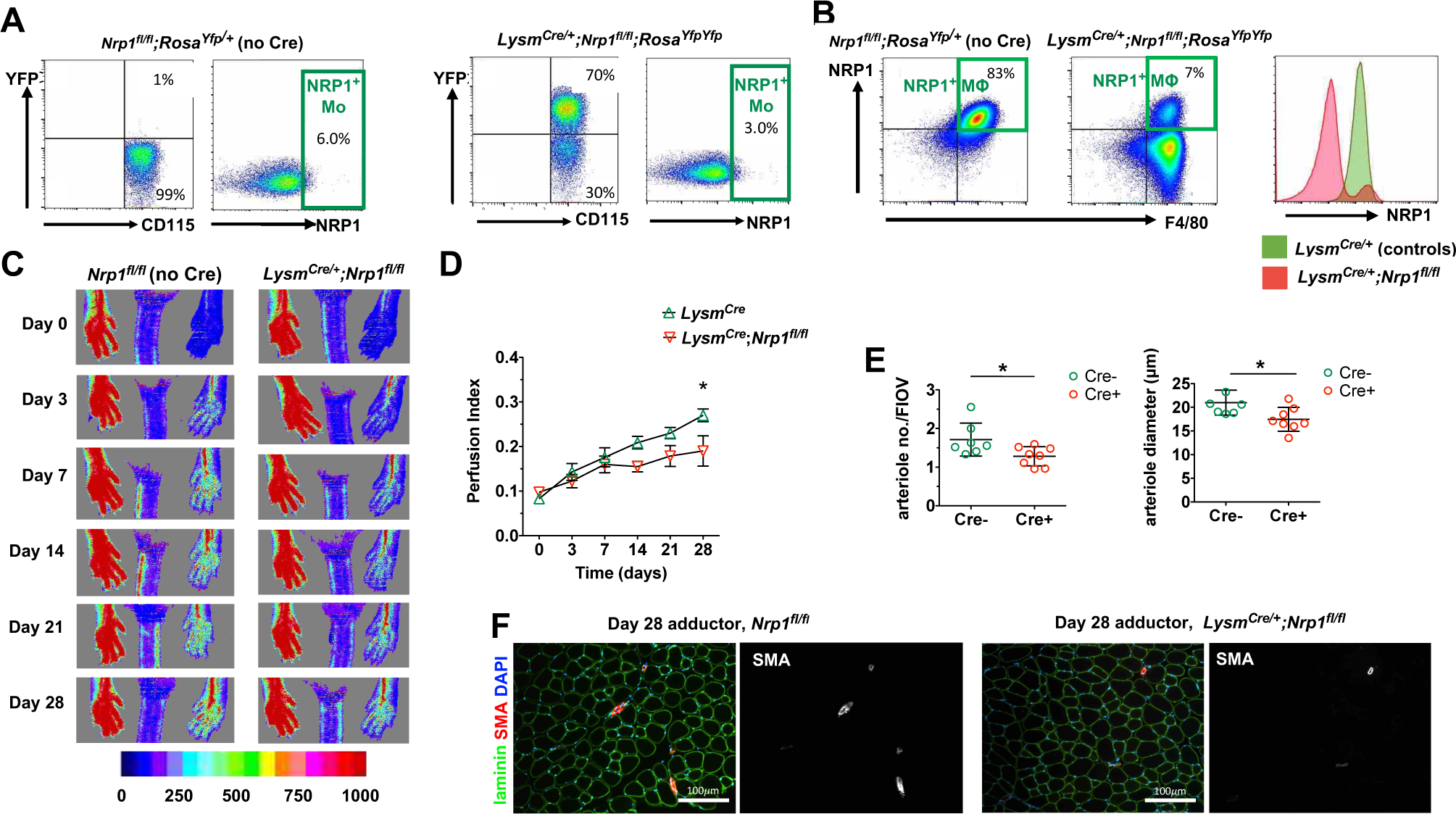
NRP1-expressing monocytes and macrophages are required for normal ischemic limb revascularization. (**A**) *Lysm^Cre^*-*mediated NRP1 targeting in monocytes and macrophages*. Flow cytometric dot plots of pre-HLI mice in circulating monocytes (**A**) and bone marrow-derived macrophages (**B**) from *Nrp1^fl/fl^;Rosa^Yfp/+^* and *Lysm^Cre/+^;Nrp1^fl/fl^;Rosa^Yfp/Yfp^* mice for the indicated markers. In (**A**), YFP analysis was used to show that *Lysm^Cre^* is active in approximately half of all circulating monocytes. In (**B**) approximately 90% reduction of NRP1 in bone marrow derived macrophages. (**C-F**) *Impaired reperfusion and arteriogenesis deficits in Lysm^Cre/+^;Nrp1^fl/fl^ mice after HLI induction*. (**C,D**) Representative flux images (**C**) and perfusion index (**D**); each data point represents the mean value ± SEM for 7 *Lysm^+/+^;Nrp1^fl/fl^* or 8 *Lysm^Cre/+^;Nrp1^fl/fl^* mice. Day 28 perfusion index: 0.19 ± 0.03 vs 0.27 ± 0.01; 2-way ANOVA P = 0.04, followed by Bonferroni’s multiple comparison test. (**E,F**) On day 28 after HLI induction, adductor muscle sections were stained for laminin and smooth muscle alpha actin (SMA) (**F**), and 27 FOV/mouse were used for quantification of arteriole density and diameter (**E**); n = 7 *Lysm^+/+^;Nrp1^fl/fl^*or 8 *Lysm^Cre/+^;Nrp1^fl/fl^* mice. Arterial density on day 28 after HLI induction (number per field of view: *Lysm^+/+^;Nrp1^fl/fl^* 1.7 ± 0.4 vs. *Lysm^Cre/+^;Nrp1^fl/fl^* 1.3 ± 0.2 (Mann-Whitney test P = 0.03). Mean arteriole diameter *Lysm^+/+^;Nrp1^fl/fl^* 21.0 ± 2.6 µm vs. *Lysm^Cre/+^;Nrp1^fl/fl^* 17.5 ± 2.5 µm (Mann-Whitney test P = 0.02). Each data point represents the value for one mouse; data are shown mean ± SD. Abbreviations: MΦ, macrophage; Mo, monocyte; * P ≤0.05.

We next asked whether supplementing NRP1-expressing macrophages could promote recovery from ischemia. Thus, we generated BMDM from wild type mice and, after sorting them according to NRP1 expression, injected them on the day of surgery into the ischemic muscle of wild type mice (**Fig. 3A; Supplementary Fig. S1C**). Doppler analysis showed that the intramuscular administration of NRP1-expressing macrophages significantly increased flow recovery following HLI when compared to the administration of NRP1-negative macrophages (**Fig. 3B,C**).

**Fig. 3.**
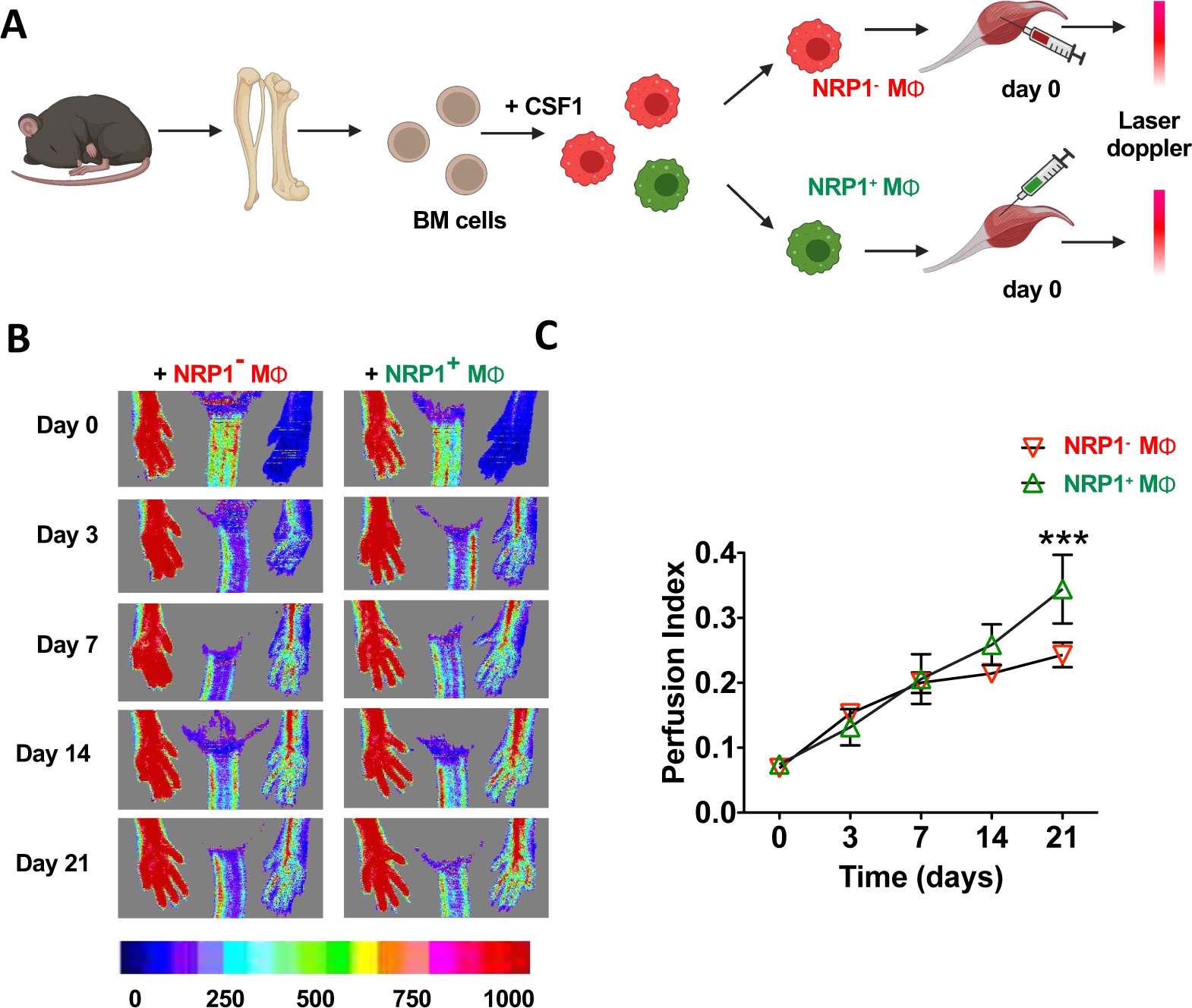
NRP1-expressing macrophages promote ischemic limb revascularization. (**A**) Schematic diagram of isolation and delivery of NRP1^+^ and NRP1^-^ bone marrow derived macrophages to ischemic limbs. (**B,C**) Ischemic muscle of wild-type mice was injected with NRP1^+^ or NRP1^-^ bone marrow-derived macrophages on the day of surgery and analysed at the indicated time points after HLI induction. Representative flux images (**B**) and perfusion index (**C**). Each data point represents the mean value ± SEM for 7 mice/group at that time point. Day 21 perfusion index: 0.34 ± 0.02 vs. 0.24 ± 0.02; 2-way ANOVA P = 0.006, followed by Bonferroni’s multiple comparison test. Abbreviations: Mχτ, macrophage; *** P ≤0.001.

In summary, loss of function and gain of function approaches show that NRP1-expressing macrophages promote arteriogenesis and flow recovery in a mouse model of CLI.

### VEGF164 induces ERK1/2 and p38 MAPK signalling in NRP1-expressing macrophages

Two main ligands for NRP1 are SEMA3A and VEGF-A^20–22^. Analysis of a publicly available transcriptomic dataset of adductor muscle in a mouse model of CLI^32^ showed that *Sema3a* was barely detected and its transcript levels were also not affected by the induction of ischemia (**Fig. 4A**). By contrast, *Vegfa* levels increased 3.5-fold at D7 and D14 after injury (**Fig. 4A**). We therefore investigated whether the NRP1-binding isoform of VEGF-A, VEGF164, induced a NRP1-dependent signalling response in bone-marrow-derived macrophages (**Fig. 4B**). As relevant readouts of VEGF164 stimulation, we used flow cytometry to examine the phosphorylation of ERK1 and ERK2 (pERK1/2), which have been implicated in arteriogenesis^38^, and the phosphorylation of P38 MAPK (pP38 MAPK), which is required for VEGF164-induced angiogenesis in a NRP1-dependent mechanism^25^. We found that VEGF164 induced significantly greater phosphorylation of both ERK1/2 and p38 MAPK in NRP1-expressing macrophages compared to NRP1-negative macrophages (**Fig. 4C**). We also examined BMDM from *Nrp1^cyto/cyto^* mice, because the NCD promotes VEGF164-induced ERK1/2 activation in endothelial cells *in vitro* and promotes arteriogenesis after HLI induction^26^. NRP1-expressing macrophages from *Nrp1^cyto/cyto^* mice showed significantly impaired VEGF164-induced ERK1/2 and p38 MAPK activation compared to NRP1-expressing macrophages from wild type mice, whereby activation levels were similarly low in *Nrp1^cyto/cyto^* and NRP1-negative BMDMs (**Fig. 4B,C**). These findings raised the possibility that NRP1 and its NCD are required to stimulate pro-arteriogenic/angiogenic macrophage signalling, a function previously attributed only to endothelial cells after HLI.

**Figure 4.**
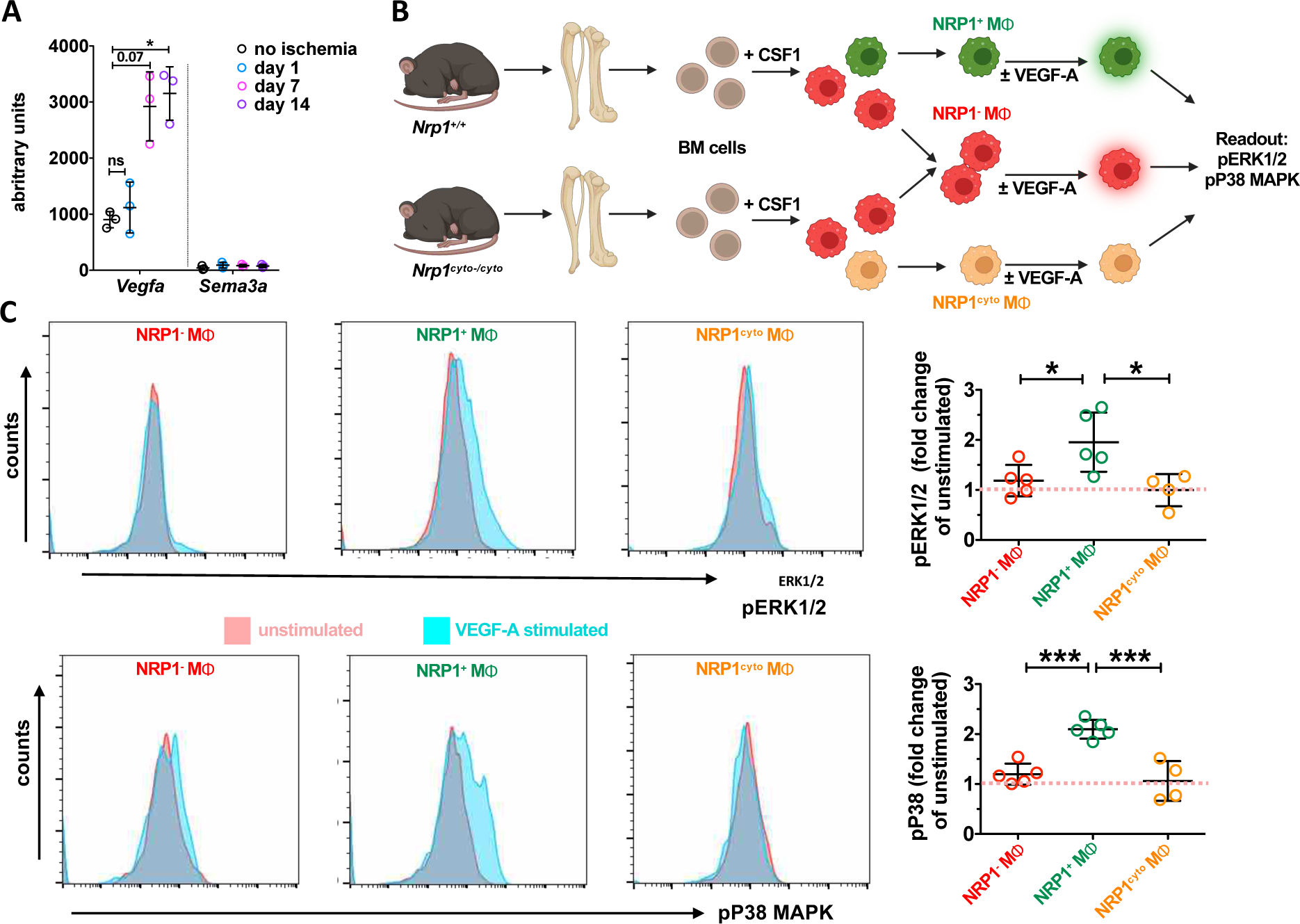
VEGF164 induces ERK1/2 and p38MAPK signalling in NRP1-expressing macrophages. (**A**) *Increased Vegfa expression in ischemic muscle*. Analysis of publicly available microarray data for *Vegfa* and *Sema3a* in ischemic wild-type adductor muscle at the indicated time-points relative to naïve muscle; n = 3 mice/group/time point; data are shown as mean ± SD; 2-way ANOVA (P = 0.002) and Dunnett’s multiple comparison test. (**B-D**) *NRP1^+^ but not NRP1^-^ or NRP1^cyto^ macrophages respond to VEGF164 stimulation.* (**B**) Schematic representation of the experimental workflow to produce bone marrow-derived NRP1^+^, NRP1^-^ and NRP1^cyto^ macrophages for measuring the response to VEGF164 stimulation *in vitro*. (**C**) Representative flow cytometric histograms showing mean fluorescent intensity as a readout of ERK1/2 and p38 MAPK phosphorylation in unstimulated versus VEGF164 stimulated macrophages, with quantification shown in (**D**) as mean fold ± SD change in VEGF164-stimulated relative to unstimulated macrophages. ERK1/2: NRP1^+^ 2.0 ± 0.6 vs. NRP1^-^ 1.2 ± 0.3 and NRP1^cyto^ 1.0 ± 0.3 (One-way ANOVA P = 0.01 and Tukey’s multiple comparison test). p38 MAPK: NRP1^+^ 2.1 ± 0.2 vs. NRP1^-^ 1.20 ± 0.2 and NRP1^cyto^ 1.1 ± 0.4 (One-way ANOVA P <0.001). Tukey’s multiple comparison test. Abbreviations: MΦ, macrophage; * P ≤0.05, *** P ≤0.001.

### Defective macrophage NRP1 signalling impairs revascularization of ischemic limbs

Corroborating prior work^26^, *Nrp1^cyto/cyto^* mice had significantly impaired reperfusion of ischemic hindlimb muscle compared to wild type mice (**Fig. 5A,B**). As previously observed with microCT analysis^26^, quantification of immunostained sections from ischemic adductor muscle showed that *Nrp1^cyto/cyto^* mice had fewer arterioles than wild-type mice (**Fig. 5C,D**). Strikingly, injecting, NRP1-expressing BMDM from wild type mice into the ischemic muscle of *Nrp1^cyto/cyto^* mice (**Fig. 5E**) rescued impaired hindlimb reperfusion (**Fig. 5F,G**). By contrast, perfusion rescue did not occur when injecting bone-marrow derived NRP1-expressing macrophages from *Nrp1^cyto/cyto^*mice into the ischemic muscle of *Nrp1^cyto/cyto^* mice (**Fig. 5E-G**). Recovery induced by NRP1-expressing macrophages from wild type mice was consistent with an increased number of arterioles compared to mice injected with NRP1-expressing macrophages from *Nrp1^cyto/cyto^* mice (**Fig. 5H,I**). These findings suggest that NRP1 expression in macrophages is functionally required for arteriogenesis in ischemic muscle.

**Fig. 5.**
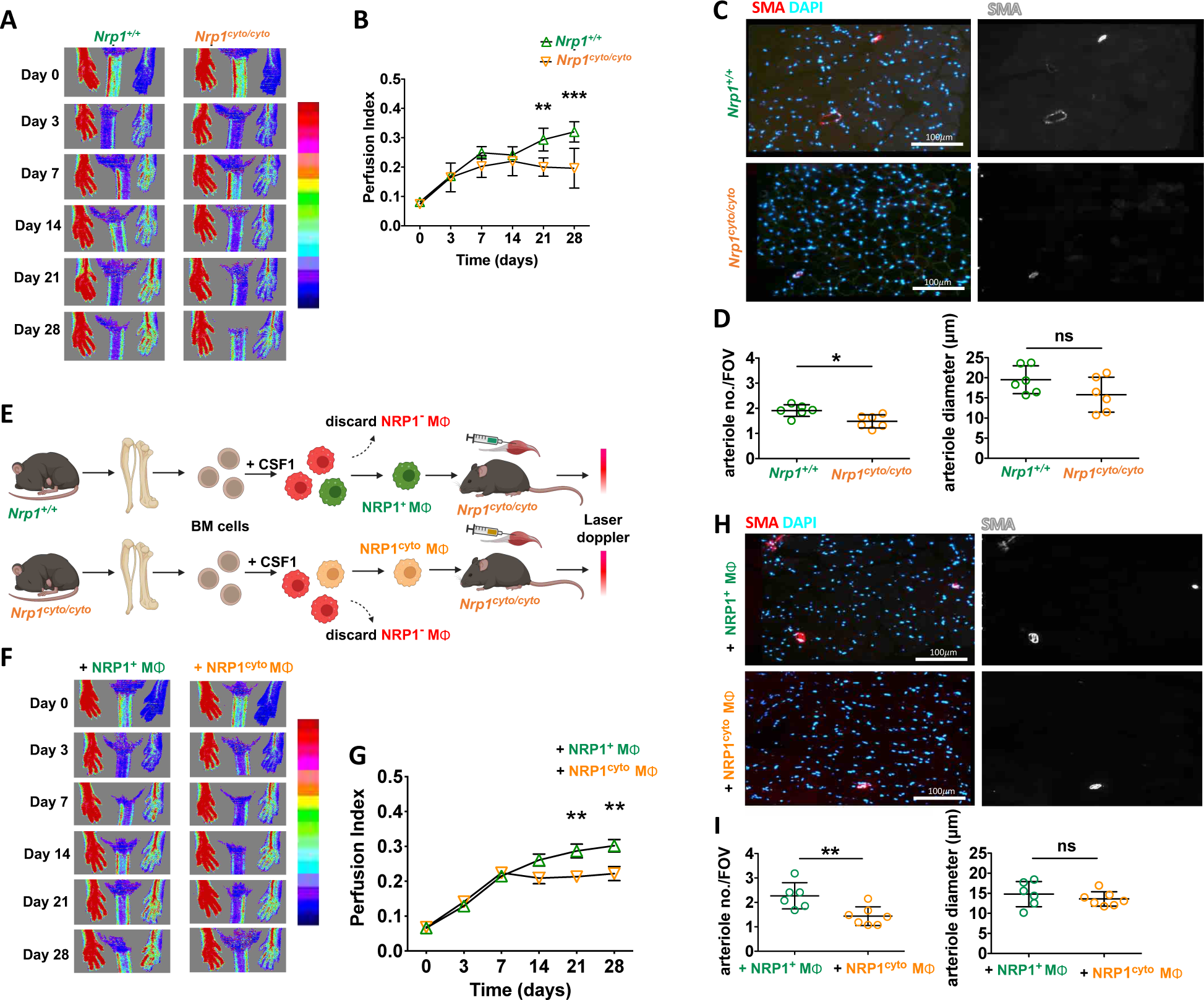
Defective macrophage NRP1 signalling impairs ischemic limb revascularization. (**A-D**) *Reperfusion and arteriogenesis deficit in Nrp1^cyto/cyto^ mice.* (**A,B)** Representative flux images (**A**) and perfusion index (**B**) of *Nrp1^+/+^* and *Nrp1^cyto/cyto^* mice at the indicated time points after HLI induction; n = 8 each. Each data point represents the mean value ± SEM from 8 mice. Day 28 perfusion index: *Nrp1^+/+^* 0.3 ± 0.03 vs. *Nrp1^cyto/cyto^* 0.2 ± 0.02, 2-way ANOVA P = 0.01, Bonferroni’s multiple comparison test. (**C,D**) On day 28, adductor muscle sections were stained for smooth muscle alpha actin (SMA) and counterstained with DAPI (**C;** scale bars 100 μm), and 27 FOV/mouse were used for quantifications (**D**); n = 6/group; each data point represents the value for one mouse. Arteriole number per field of view, mean ± SD: *Nrp1^+/+^* 1.9 ± 0.2 vs. *Nrp1^cyto/cyto^* 1.5 ± 0.3, Mann-Whitney test, P = 0.02. Arteriole diameter: *Nrp1^+/+^* 19.5 ± 3.5 µm vs. *Nrp1^cyto/cyto^* 15.8 ± 4.4 µm, Mann-Whitney test, P = 0.24. (**E-I**) *NRP1^+^ macrophages rescue arteriogenesis deficits of Nrp1^cyto/cyto^ mice. Nrp1^cyto/cyto^* mice were injected with bone marrow-derived NRP1-expressing macrophages from wild-type (NRP1^+^ MΦ) or *Nrp1^cyto/cyto^* (NRP1^cyto^ MΦ) mice on the day of surgery. (**E**) Schematic diagram of the work flow. (**F,G**) Representative flux images (**F**) and perfusion index (**G**) at the indicated time points after HLI induction. Each data point represents the mean value ± SEM for 10 mice at that time point. Day 28 perfusion index: NRP1^+^ MΦ 0.3 ± 0.02 vs. NRP1^cyto^ MΦ 0.2 ± 0.02; 2-way ANOVA P = 0.02, Bonferroni’s multiple comparison test. (**H,I**) On day 28, adductor muscle sections were stained for smooth muscle alpha actin (SMA) and counterstained with DAPI (**H;** scale bars 100 µm), and 27 FOV/mouse were used for quantification (**I**) of arteriole density and diameter; NRP1^+^ MΦ n= 6 mice, NRP1^cyto^ MΦ n = 7 mice. Arteriole number per field of view, mean ± SD: NRP1^+^ MΦ 2.3 ± 0.5 vs. NRP1^cyto^ MΦ1.4 ± 0.4, Mann-Whitney test, P = 0.008. Arteriole diameter: NRP1^+^ MΦ 14.8 ± 3.1 µm vs. NRP1^cyto^ MΦ13.6 ± 1.8 µm, Mann-Whitney test, P = 0.45. Each data point represents the value for one mouse; data are shown mean ± SD. Abbreviations: MΦ, macrophage; ns, not significant (P >0.05), *<0.05, ** P <0.01, ***<0.001

### NRP1-expressing monocytes from CLI patients stimulate SMC proliferation and EC tubulogenesis

We next sought to determine the relevance of our findings in the mouse HLI model for CLI patients In keeping with our observation in the mous, peripheral blood from CLI patients contained an increased proportion of NRP1-expressing monocytes compared with peripheral blood from age-matched, healthy control subjects (**Fig. 6A,B**; Supplementary **Fig. S1D**). Human circulating monocytes have been classified as classical (CD14^++^ CD16^-^) and tissue remodelling subpopulations which include intermediate (CD14^++^ CD16^+^) and non-classical (CD14^+^ CD16^++^) cells^39^ (**Fig. 6C,D**). The greatest proportion of NRP1-expressing monocytes were found in the intermediate population (**Fig. 6C,E**). This intermediate population also contained a high proportion of cells expressing CD163 (**Fig. 6F**), a marker of monocytes with tissue remodelling properties^40^, and contained monocytes expressing VEGFR2, albeit their enrichment in the intermediate population compared to the other two populations was not significant (**Fig. 6G**).

**Fig 6.**
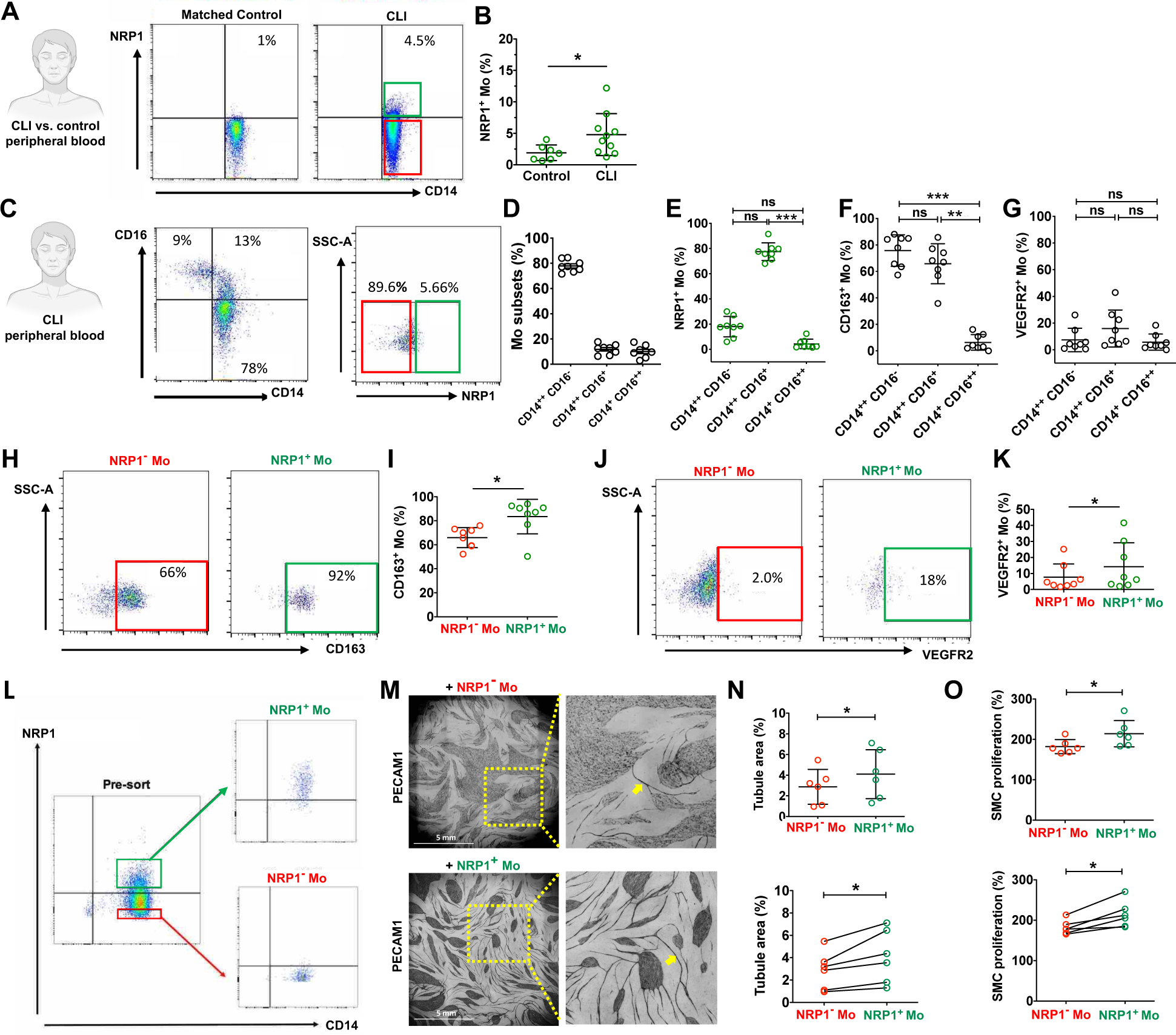
NRP1-expressing monocytes from CLI patients stimulate vascular endothelial tubulogenesis and SMC proliferation. (**A,B**) *NRP1^+^ circulating monocytes are increased in CLI patients compared to controls*. (**A**) Representative flow cytometric dot-plots from one CLI patient and a matched control. (**B**) quantification of CD14^+^ NRP1^+^ monocytes in all monocytes (Lin^-^ CD14^+^) in 10 CLI patients versus 7 age-matched controls: control 1.9 ± 0.5% vs. CLI 4.8 ± 1.1%; Mann-Whitney test P = 0.03. (**C-E**) *NRP1^+^ circulating monocytes of CLI patients are enriched in the intermediate population.* (**C**) Representative flow cytometric dot-plots of monocytes analysed for CD14 vs CD16 and NRP1 from whole monocyte population. (**D,E**) Proportion of NRP1^+^ monocytes in the classical (CD14^++^CD16^-^), intermediate (CD14^++^CD16^+^) and non-classical (CD14^+^CD16^++^) populations; n = 8 CLI patients. (**D**) CD14^++^CD16^-^ 77.9 ± 4.7%, CD14^++^CD16^+^ 11.8 ± 3.9%, CD14^+^CD16^++^ 10.1 ± 4.8%. (**E**) CD14^++^CD16^-^ NRP1^+^ 18.0 ± 8.1% vs. CD14^++^CD16^+^NRP1^+^ 77.5 ± 7.0% and CD14^+^CD16^++^NRP1^+^ 4.3 ± 3.8%; Kruskal-Wallis test P<0.0001 followed by Dunn’s multiple comparison test. n = 8 CLI patients. Each data point represents the value from one patient. (**F**) CD14^++^CD16^-^CD163^+^ 75.8 ± 11.9% vs CD14^++^CD16^+^CD163^+^ 65.8 ± 15.1% and CD14^+^CD16^++^CD163^+^ 6.4 ± 5.9%; Kruskal-Wallis test P=0.0009 followed by Dunn’s multiple comparison test, n = 8. (**G**) CD14^++^CD16^-^VEGFR2^+^ 7.4 ± 8.8 vs CD14^++^CD16^+^VEGFR2^+^ 16.0 ± 13.8 and CD14^+^CD16^++^VEGFR2^+^ 5.7 ± 6.0; ; data are shown mean ± SD; Kruskal-Wallis test P=0.0003 followed by Dunn’s multiple comparison test n = 8. (**H-K**) *VEGFR2 and CD163 enrichment in NRP1^+^ vs. NRP1^-^ CLI circulating monocytes.* Representative flow cytometric dot-plots (**H,J**) and quantification (**I,K**); n = 8 CLI patients. Each data point represents the value from one patient. (**I**) Proportion of NRP1^+^ and NRP1^-^ monocytes that express CD163: NRP1^-^ 65.9 ± 8.3% vs. NRP1^+^ 83.5 ± 14.4%; Wilcoxon matched-pairs signed rank test, P = 0.02. (**K**) Proportion of NRP1^+^ and NRP1^-^ monocytes that express VEGFR2: NRP1^-^ 7.7 ± 8.3%, vs. NRP1^+^ 14.3 ± 14.9%; data are shown mean ± SD; Wilcoxon matched-pairs signed rank test P = 0.02. (**L-N**) *NRP1^+^ circulating monocytes from CLI patients promote angiogenesis in vitro.* NRP1^+^ and NRP1^-^ monocytes were isolated from the blood of patients and used as matched pairs in endothelial/fibroblast co-culture assay. (**L**) Representative flow cytometric dot-plots of fluorescence activated cell sorting and (**M**) PECAM1-stained cultures after incubation with CLI monocytes for 14 days. (**N**) Quantification of tubule area as an indicator of angiogenic sprouting; n = 6 matched NRP1^+^ vs. NRP1^-^ monocyte pairs from CLI patients. Data are shown as mean ± SD (top graph) and as matched pairs (bottom graph). Each data point represents the NRP1^+^ vs. NRP1^-^ monocyte population from on CLI patient. NRP1^-^ 2.9 ± 1.7% vs. NRP1^+^ 4.1 ± 2.4%, Wilcoxon matched-pairs signed rank test, P = 0.03. (**O**) Conditioned media from *NRP1^+^ circulating monocytes from CLI patients promote SMC proliferation in vitro.* SMC proliferation was quantified as relative change; n = 6 matched NRP1^+^ vs. NRP1^-^ monocyte pairs from CLI patients. Data are shown as mean ± SD (top graph) and as matched pairs (bottom graph). Each data point represents the NRP1^+^ vs. NRP1^-^ monocyte population from on CLI patient. NRP1^-^ 182.2 ± 17.4% vs. NRP1^+^ 214.2 ± 32.5%, Wilcoxon matched-pairs signed rank test, P = 0.03. Abbreviations: CLI, critical limb ischemia; Mo, monocyte; ns, not significant (P >0.05), *<0.05, ***<0.001.

We next examined directly whether monocytes expressing CD163 and VEGFR2 were enriched in the NRP1-expressing monocyte population of CLI patients (**Fig. 6H-K**). We found that the NRP1-expressing monocyte population was significantly enriched in cells expressing CD163 (**Fig. 6H,I**) and VEGFR2 (**Fig. 6J,K**) when compared to the population of NRP1-negative monocytes. These findings suggest that human NRP1-expressing monocytes have a tissue remodelling phenotype, similar to NRP1-expressing monocytes and macrophages from mice after HLI induction (see **Fig. 1**). Agreeing with this idea, NRP1-expressing monocyte isolated from the blood of CLI patients (**Fig. 6L**) stimulated endothelial tubule formation (**Fig. 6M,N**) and SMC proliferation (**Fig. 6O**) more than NRP1-negative monocytes isolated from the same patients.

Taken together with our mechanistic findings using the mouse HLI model, these observations suggest that NRP1-expressing monocytes increase in response to ischemia and that NRP1-expressing macrophages stimulate ischemic neovascularization, both endogenously and when delivered therapeutically, to enhance revascularisation and accelerate restoration of perfusion (working model in **Fig. 7**).

**Figure 7.**
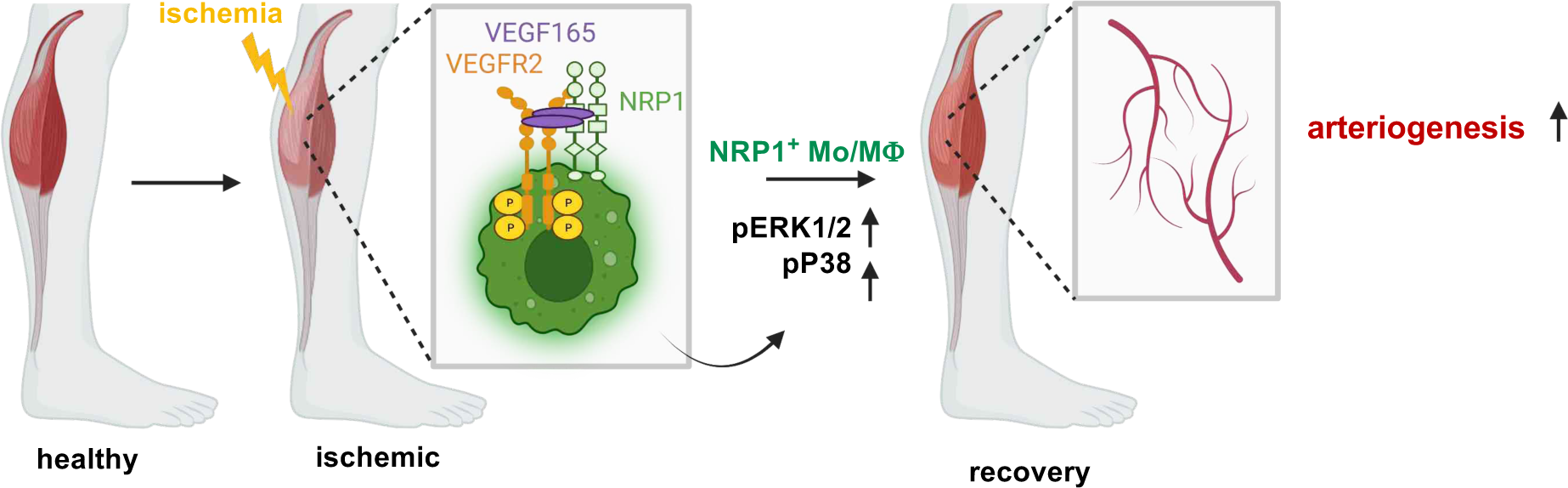
Working model. Ischemia (yellow bolt) induces expression of VEGF165 (purple symbols). VEGF165 signalling in NRP1-expressing monocytes and macrophages (green cells), indicated by phosphorylation of ERK1/2 and p38 MAPK, promotes revascularisation of the ischemic limb.

## Discussion

In the present study, we have found that the proportion of circulating monocytes that express NRP1 was raised in mice after from D3 in a mouse mode of CLI and in patients with CLI compared with healthy controls. Using gene expression studies and flow cytometry in the mouse model, we could further show that the proportion of macrophages expressing NRP1 also increased in ischemic compared to non-ischemic adductor muscle from D3 after induction of ischemia. The increased proportion of NRP1-expressing macrophages observed by flow cytometry was reflected in an upregulation of NRP1 and macrophage marker gene expression in the ischemic muscle and coincided both with the upregulation of endogenous VEGFA expression, the onset of arterial marker expression and improved blood flow recovery (**Figs. 1** and **4**). Together, these findings raise the possibility that NRP1-expressing macrophages increase in number in the ischemic muscle due to the VEGF165-induced recruitment or expansion. In agreement with this idea, adenoviral VEGF165 over-expression in mouse tumours or in normal mouse muscle also recruits monocytes via NRP1^3041^. Although the alternative NRP1 ligand SEMA3A can also induce the recruitment of NRP1-expressing macrophages to tumours^29^, we found that its expression is very low in the uninjured muscle and not upregulated after induction of HLI (**Fig. 4**). Therefore, VEGF164 rather than SEMA3A appears to be the relevant factor for NRP1-expressing macrophages in ischemic murine tissue.

In the mouse CLI model, a greater proportion of NRP1-expressing macrophages co-expressed MRC1 when compared to NRP1-negative macrophages, and MRC1 expression levels were accordingly increased in the ischemic adductor muscle (**Fig. 1**). The proportion of NRP1-expressing monocytes that expressed MRC1 was also increased in the circulation (**Fig. 1**). These findings suggest that NRP1-expressing monocytes and macrophages are skewed towards the so-called ‘M2-polarisation’ phenotype, which is considered to reflect a tissue remodelling phenotype that promotes repair and neovascularization^42^ ^43^. In agreement with this idea, and suggesting clinical relevance, the proportion of CD163-expressing monocytes with tissue remodelling properties^40^ was higher in the pool of NRP1-positive compared to NRP1-negative monocytes also in the blood from CLI patients (**Fig. 6**). Further, NRP1-expressing monocytes from the blood of CLI patients were enriched within the CD14^++^ CD16^+^ population (**Fig. 6**), which has been proposed to be the pro-angio/arteriogenic monocyte subset^44–46^.

NRP1-expressing myeloid cells have previously been reported to promote choroidal neovascularization^47^, and NRP1 is also required for myeloid recruitment to non-ischemic muscle overexpressing adenoviral VEGF-A^30^. To determine whether myeloid NRP1 is functionally important for recovery from ischemia, we used *Lysm^Cre/+^;Nrp1^fl/fl^* mice, which had approximately 50% fewer NRP1-expressing monocytes and macrophages. Reperfusion recovered in these mice less effectively than in control mice, and this was explained by reduced muscle arteriole number and density compared to controls (**Fig. 2**). Together, these data suggest that NRP1-expressing macrophages promote revascularization of ischemic tissue by promoting arteriogenesis. In agreement, NRP1-expressing monocytes isolated from CLI patients had greater angiogenic potency and increased SMC proliferation more than NRP1-negative monocytes (**Fig. 6**). These data agree with previous studies using adenoviral VEGF165 over-expression, which showed that NRP1-expressing monocytes promote arteriogenesis possibly by secreting factors such as PDGF-B^29^, but extend prior work by demonstrating that VEGF165 activates NRP1 signalling in macrophages in the setting of muscle ischemia.

Although NRP1-expressing macrophages and monocytes promote arteriogenesis, NRP1-expressing monocytes persist in the circulation of both in a CLI mouse model and in patients with CLI, suggesting that their recruitment into the ischemic muscle is incomplete. We therefore examined whether intra-muscular delivery of these cells would accelerate re-vascularization. Indeed, delivery of NRP1-expressing macrophages directly into the adductor muscle immediately after surgery in the CLI mouse model improved revascularization from D7 onwards, with the greatest difference on D21, the latest time point examined. Intra-muscular injection was chosen as the mode of cell delivery in our study, because meta-analyses of studies investigating cell therapies for limb ischemia have shown that intra-muscular injection is more effective than intra-arterial delivery^4^ ^48^. The inefficiency of intra-arterial injection may be explained by cells not reaching the areas of ischemia due to fewer patent vessels in the ischemic muscle. By contrast, cells delivered locally to the ischemic muscle can act through paracrine mechanisms to promote neovascularization^29^, and agreeing with our finding found that NRP1-expressing monocytes promoted endothelial tube formation and SMC proliferation (**Fig. 6**). Notably, there was no difference in perfusion between ischemic hindlimbs injected with NRP1 positive and NRP1 negative macrophages up to 7 days after HLI **(Fig. 3**). This suggests that the macrophage-mediated effect does not take hold immediately, perhaps because the injected cells need to adapt to the ischemic muscle environment and activate the vasculature, which itself has to undergo vascular remodelling and vascular smooth muscle induction for clinically meaningful neovascularization.

A subset of peripheral monocytes expresses VEGFR2 and are deemed proangiogenic^49^ ^50^. We identified VEGFR2-expressing monocytes amongst the NRP1-expressing monocytes compared to NRP1-negative monocytes (**Fig. 6**). This finding raises the possibility that VEGF165 stimulation could induce complex formation of NRP1 and VEGFR2 in monocytes, as previously shown for endothelial cells. In agreement, we observed that the endothelial VEGF165/NRP1/VEGFR2 downstream target ERK1/2 was activated in NRP1-expessing macrophages (**Fig. 4**), as previously shown in cultured endothelial cells, where NRP1 knockout decreases VEGF165-induced ERK1/2 activation^26^. In endothelial cells, NRP1’s NCD is thought to promote pro-arteriogenic ERK1/2 signalling from VEGF165/NRP1/VEGFR2 complexes by promoting NRP1/GIPC1-mediated VEGFR2 endocytosis^27^ ^28^. To investigate the importance of NRP1-mediated VEGF165 downstream signalling in macrophages, we used NRP1-expressing BMDMs isolated from *Nrp1^cyto/cyto^* mice; in these mice, the NCD is lacking, but NRP1 protein expression is retained^51^, thereby allowing us to isolate NRP1-expressing macrophages from these mice, unlike from *Lysm;Nrp1^fl/fl^*mice lacking NRP1 altogether in the myeloid lineage. Comparing NRP1-expressing macrophages lacking or retaining the NCD showed that VEGF165-stimulated ERK1/2 signalling occurs in macrophages and requires the NCD, as described for endothelial cells. Furthermore, we observed an NCD dependence of P38 MAPK activation, which was previously implicated in NRP1-dependent angiogenesis^25^ but is also involved in the alternative activation of macrophages towards the M2-like tissue remodelling phenotype^52^.

NCD-dependent signalling is clearly important for stimulation of neovascularization in the ischemic muscle, because *Nrp1^cyto/cyto^* mice had impaired recovery of perfusion in the ischemic limb, which was associated with a decreased arteriole density (**Fig. 5**)^26^. As *Nrp1^cyto/cyto^* mice lack the NCD in all cells, including endothelial cells, they do not allow distinguishing whether VEGF165-NRP1 signalling in monocytes and macrophages rather than in endothelial cells contributes to recovery from ischemia. We therefore injected NRP1-expressing macrophages from wild type mice to rescue impaired recovery in *Nrp1^cyto/cyto^* mice. Longitudinal assessment of perfusion showed improved reperfusion of the ischemic limb from D7 onwards, and greater arteriole density on D28, similar to experiments in which wild type NRP1-expressing macrophages were injected into wild type mice. This experiment suggests that NCD-dependent signalling for recovery from ischemia takes place in NRP1-expressing macrophages.

We conclude that NRP1-expressing macrophages promote recovery from muscle ischemia by stimulating intramuscular arteriogenesis, their number is usually rate-limiting, targeted, therapeutic delivery of larger numbers monocyte-derived, NRP1-expressing macrophages might, therefore, enhance vascular remodelling for improved recovery of perfusion. Accordingly, the autologous delivery of peripheral blood-derived, NRP1-expressing monocytes or their functional equivalents should be investigated as a potential treatment in patients with intractable limb or myocardial ischemia.

## Acknowledgments

We thank the National Institute of Health Research (NIHR) Guy’s and St Thomas’ Biomedical Research Centre (BRC) and the Nikon Imaging Centre at King’s College London for providing equipment and technical assistance. We thank Professor Harry Mellor for advice on the tubulogenesis assay and Dr Richard Siow for providing human umbilical vein endothelial cells (HUVECs). We thank the UCL Institute of Ophthalmology Biological Resource Unit for mouse husbandry.

## Sources of funding

This work was supported by the British Society of Endovascular Therapy and Royal College of Surgeons of England fellowship to JC, British Heart Foundation grant to BM (FS/17/24/32596) and Wellcome Investigator Award to CR (205099/Z/16/Z).

## Disclosures

None.

## Methods

### Animals

All animal procedures were performed according to UK Home Office and local Animal Welfare and Review Body (AWERB) guidelines. We combined *Nrp1^fl/fl^;Rosa^yfp/yfp^ with Lysm^Cre^* (*Lyz2^Cre^*) to obtain *Lysm^Cre/+^;Nrp1^fl/fl^* mutants and *Lysm^+/+^;Nrp1^fl/fl^*controls^37,53,54^. We also used *Nrp1^cyto/cyto^*mice and *Nrp1^+/+^* (WT) controls^51,55^. All mouse strains were maintained on a C57/Bl6J background.

### Murine hindlimb ischemia

Hindlimb ischemia was induced by excising a section of the femoral artery, as previously described^31^. Male mice aged between 8 to 12 weeks were used for the model. Ischemia was confirmed by laser Doppler (Moor) scanning. In some experiments, 1 x 10^6^ cells in PBS or vehicle control (PBS) were injected into the ischemic hindlimb adductor muscle on day 0. Longitudinal blood perfusion measurements were obtained by laser Doppler scanning of ischemic and contralateral limbs at days 0, 3, 7, 14, 21 and 28 post HLI. Moor image analysis software was used where ROI was created and the median value of the ischemic hindlimb was divided by the control to determine the perfusion index.

### Histological muscle analysis

Ipsilateral murine adductor muscles were harvested and then fixed in 1 ml of 4% formaldehyde for 30 min at RT followed by 24 h each in 15%, 30% and then 40% sucrose PBS. The muscle was then mounted on cork with optimum cutting temperature (OCT) compound and snap frozen in isopentane equilibrated in liquid nitrogen. Muscle specimens were cut into 10 μm slices and stored at -80°C. Slides were air dried at RT for 10 min and then fixed in acetone at -20°C for 10 min. A PAP pen (Abcam) was used to draw around the tissue and blocking reagent (2.5ml FCS, 0.5 g BSA, 50 μl TritonX-100 in 50 ml PBS) was added for 1 h at RT. Blocking reagent was tapped off and antibodies were added for 1 h at RT. Arteriole diameter and number were measured on sections stained with the following antibodies: anti-actin, a-smooth muscle-cy3 antibody (Sigma-Aldrich) and Dylight 488-conjugated anti-laminin (Novus Biologicals). Then the slides were washed 3 times for 5 minutes in 500 ml PBS containing 250 μl Tween 20. 1 drop of DAPI-Vectashield mountant (Vector laboratories) was added, a coverslip placed on top and sealed with varnish. Three fields of view/section for three sections at three different muscle levels were analyzed, then average value used. Fluorescent staining was assessed with a NikonTi Eclipse microscope using NIS-Elements BR microscopy software.

### Patient recruitment

Patients with critical limb ischemia (rest pain ± tissue loss) and healthy age- and sex-matched controls were recruited into this study with research ethics committee approval 10/H0804/67. Controls were recruited from volunteers who were admitted for elective surgery. Patients with active malignancy or those taking steroids were excluded.

### Flow Cytometry

Cell phenotypes were assessed using a MACSQuant (Miltenyi Biotec, UK) or AttuneNxT (Thermo Scientific, UK) flow cytometer. Isotype controls or fluorescence minus one (FMO) controls were used, and the live/dead marker 7-aminoactinomycin D (7-AAD, PerCP) (Miltenyi Biotec) was added to all samples 10 min before analysis (10 μl to 1 ml of cell suspension).

#### Analysis of NRP1-expressing monocytes in murine peripheral blood

Mice were anesthetized using isoflurane (Abbott, UK) and blood was collected through the heart using a 1 ml insulin syringe with a 29G needle before HLI as well as on days 3, 7, 14 and 21 post HLI. 400 μl blood was added to 20 ml of red blood cell lysis buffer (50 ml deionized water, 450 mg ammonium chloride, 50 mg sodium hydroxide, 18 mg EDTA, pH 7.4) for 30 min in RT. Then 30 ml of PBS containing 0.5% BSA and 2 mM EDTA was added and the suspension centrifuged at 400 g for 5 min at 4°C. The supernatant was discarded, and 20 ml of phosphate buffered saline (PBS) containing 0.5% bovine serum albumin (BSA) and 2mM EDTA (termed FACS butter) was added to the pellet. The cell suspension was centrifuged again at 400g for 5 min at 4°C and the supernatant was discarded again. The cells were resuspended in 200 μl of PBS containing 0.5% BSA and 2 mM EDTA and incubated with 15 μl of mouse FcR blocking reagent (Miltenyi Biotec) on ice for 15 min. Subsequently, antibodies were added for 1 h on ice in the dark (Table 1). After incubation, the cells were washed with 2 ml of FACS buffer and resuspended in 400 μl FACS buffer for analysis using the gating strategy in Supplementary Fig. S1B.

#### Quantification of NRP1-expressing macrophages in murine muscle

The adductor (medial thigh) muscle was harvested pre-HLI and from the ischemic limb at 3 and 7 days post-HLI. Muscle was crushed with a 20 ml syringe plunger against a 70 μm cell strainer (Corning) into a Petri dish containing PBS. The cell suspensions were individually transferred into 50 ml Falcon tubes and centrifuged at 400 g for 5 min at 4°C. Red blood cells were lysed as described above. Subsequently, antibodies were added (Table 1) and labelled cells analyzed using the gating strategy in Supplementary Fig. S1A.

#### Analysis of NRP1-expressing monocytes in human peripheral blood

Peripheral venous blood from patients was collected into ethylenediaminetetraacetic acid (EDTA) anti-coagulated tubes (BD Vacutainer, UK). 400 μl blood was added to 10 ml of 1x lysis solution (BD Pharm Lyse, BD Biosciences) for 30 min at room temperature (RT) before 30 ml of FACS buffer was added. The samples were centrifuged at 400 g for 5 min at 4°C, the supernatant was discarded, and 20 ml of flow cytometry buffer was added to the pellet before the sample was centrifuged again at 400 g for 5 min at 4°C. The supernatant was discarded again, and the cells were resuspended in 200 μl of flow cytometry buffer and incubated with 15 μl of human FcR blocking reagent (Miltenyi Biotec) on ice for 15 min. Subsequently, antibodies were added for 1 h on ice in the dark (Table 1). After incubation, the cells were washed with 2 ml of flow cytometry buffer and resuspended in 400 μl of flow cytometry buffer for analysis. Monocytes were gated as shown in Supplementary Fig. S1D.

#### Generation and isolation of bone marrow derived macrophages (BMDMs)

Mice were culled by cervical dislocation. Hindlimbs were amputated and dissected down to the bone. The bones were divided at the joint to separate the femur and tibia. In the cell culture hood, the marrow was flushed with a 23G needle and 20 ml of sterile PBS into a 50 ml Falcon tube. The cell suspension was filtered through a 70 μm cell strainer (Corning) and centrifuged for 5 min at 1400 rpm at RT. The supernatant was discarded and 5 ml of red blood cell lysis buffer (as above) was added for 5 min at RT and then 10 ml RPMI-1640 (Thermofisher) supplemented with antibiotic/antimycotic, L-glutamine and 10% FCS (termed macrophage media). After centrifugation for 5 min at 1400 rpm at RT, the supernatant was discarded, the cells were resuspended in macrophage media containing 50 ng/ml CSF1 (M-CSF) and cultured at 37°C in 5% CO2 with media change every 2-3 days. Media was aspirated and then discarded from confluent T75 x1 and T175 x2 culture flasks, and BMDMs were scraped with cell scrapers (Corning) in PBS. Cells were centrifuged at 400 g for 5 min at 4°C, the supernatant was discarded and the cells were resuspended in 200 μl of flow cytometry buffer. Cells were incubated with 15 μl of mouse FcR blocking reagent (Miltenyi Biotec) on ice for 15 min. PE-conjugated anti-NRP1 (Miltenyi Biotec) and PE-Cy7-conjugated anti-F4/80 (Miltenyi Biotec) were added for 1 h on ice in the dark. Then, the cells were washed with 2 ml of buffer, centrifuged at 400 g for 5 min at 4°C and resuspended in 1 ml of flow cytometry buffer. Cells were sorted using the FACS Aria II into NRP1^+^ and NRP1^-^ macrophage populations.

#### Fluorescence activated cell sorting of human circulating monocytes

Peripheral venous blood from patients with CLI and controls was collected into ethylenediaminetetraacetic acid (EDTA) anti-coagulated tubes (BD Vacutainer, UK). 17.5 ml of whole blood was added to sterile 50 ml falcon tube and diluted 1:1 with cold sterile DMEM. This was carefully layered on top of 15 ml of Ficoll-Paque Premium (Cytiva), minimizing mixing at the Ficoll/blood interface, and centrifuged at 1400 rpm for 30 min at 4°C (with no brake). The buffy layer was collected and transferred to sterile 50 ml Falcon tube and diluted 1:1 with PBS containing 0.5% BSA and 2 mM EDTA. This cell suspension was centrifuged at 1100 rpm for 5 min at 4°C (with brakes on). The supernatant was discarded and the tube vortexed. Remaining RBCs were lysed using 1x RBC lysis buffer (BD Pharm Lyse, BD Biosciences) at RT for 5 min and the suspensions was then diluted with DPBS containing 0.5% BSA. This suspension was centrifuged again at 1400 rpm for 5 min and resuspended in 500 μl of PBS containing 0.5% BSA. FcR human blocking reagent (Miltenyi Biotec) and CD14 MicroBeads (Miltenyi Biotec) were added to the isolated peripheral blood mononuclear cells (PBMC) and incubated for 15 min at 2-8°C before the cells were washed using PBS containing 0.5% BSA and resuspended again. Magnetic separation was achieved with an LS column (Miltenyi Biotec) as per manufacturer’s instruction. CD14^+^ cells were stained using PE-conjugated anti-NRP1 (Miltenyi Biotec) and FITC-conjugated anti-CD14 (Miltenyi Biotec) antibodies (see table 2), and cells were sorted into NRP1^+^ and NRP1^-^ monocyte populations using the FACS Aria II (BD Biosciences).

### Murine phosflow

NRP1^+^ and NRP1^-^ macrophages were generated from BMDMs from WT and *Nrp1^cyto/cyto^* mice using cell sorting as described above. The cells were counted and divided into three similarly sized groups. 10 µl of Halt phosphatase inhibitor cocktail (Thermo Scientific) was added, then the cells were stimulated with 50 ng/ml VEGF164 (R&D Systems) or not stimulated for 5min in 37°C water bath. Cells were fixed with Fix buffer I (BD Bioscience), then permeabilized with Perm buffer III (BD Biosciences) as per product data sheets, then stained with 20 µl of either pERK1/2 or p38 MAPK antibodies (BD Biosciences) in 100 µl and incubated for 30 min on ice in the dark. Analysis was carried out by flow cytometry, where median fluorescence intensity (MFI) was obtained. The fold change was calculated from matched VEGF164-stimulated relative to unstimulated cells.

### HUVEC/fibroblast co-culture angiogenesis assay

Protocols were adapted from a published method^56^. Normal human dermal fibroblasts (NHDF; PromoCell, UK, up to passage 12) were cultured in DMEM (Sigma-Aldrich) supplemented with 10% FCS, 100 U/ml penicillin, 100 μg/ml streptomycin and 250 ng/ml antimycotic and maintained at 37°C in a humidified atmosphere of 5% CO_2_. NHDF were suspended at 4.8 x 10^4^ cells/ml and plated in 24-well plates (500 μl/well) and cultured for 4 days to allow a monolayer to form. HUVECs (passage 2-4; isolated from donors were provided by Dr Siow) cultured in endothelial cell growth medium (EGM; PromoCell) were resuspended at 3 x 10^4^ cells/ml and plated on top of the NHDF (500 μl/well for 24 well plates). HUVECs were allowed to attach to the fibroblasts for 6 hours at 37°C before cell-sorted monocytes were added at 3 x 10^4^ cells/ml (500 μl/well) with macrophage media (RPMI 1640 media containing 5 ml L-glutamine, 5 ml antibiotic, antimycotic, 50 ml FCS and 0.5 μl/ml CSF1). Plates were incubated at 37°C in a humidified atmosphere with 5% CO2 for 14 days, with EGM every 2-3 days. After 14 days, cells were fixed in ice cold 70% ethanol for 30 min (500 μl/well) then washed with PBS. Endogenous peroxidase activity was blocked by incubating cells with 0.3% hydrogen peroxide in methanol (Sigma Aldrich, UK) for 15 mins at room temperature (RT) followed by a PBS wash. Cells were then incubated for 1 h at 37°C with mouse anti-human PECAM1 (R&D Systems), which was diluted 1:1000 in 1% BSA/PBS using 300 μl/well. After PBS washes, the cells were incubated for 1 h at 37°C with rabbit anti-mouse IgG alkaline phosphatase conjugate, diluted 1:500 in 1% BSA/PBS using 300 μl/well, which was again followed by PBS washes. 500 μl BCIP/NBT solution (Sigma, 1 tablet in 10 ml of water, filtered with a 0.2 μm filter) was added to each well and incubated at 37°C for 15 min. Cells were washed with distilled water. Images were taken on NikonTi Eclipse microscope using NIS-Elements BR software and analyzed with ImageJ version 2.0.0-rc-68/1.52f (NIH Bethesda).

### Smooth muscle cell proliferation assay

Human aortic smooth muscle cells (SMCs; Lonza, UK) were seeded into a 96-well plate at a density of 1×10^3^/well and incubated for 24 h with 100 µl/well of Iscove’s Modified Dulbecco’s Media (IMDM) media, previously conditioned by NRP1-expressing or NRP1-negative monocytes at 37°C in a humidified atmosphere with 5% CO2. After adding XTT (sodium 3’-[1- (phenylaminocarbonyl)- 3,4- tetrazolium]-bis (4-methoxy6-nitro) benzene sulfonic acid hydrate) (1 mg/ml) to cells, it was incubated for 4 h at 37°C, after which absorbance was measured at 450 nm using a plate reader (Filter Max F5, Molecular Devices) with a reference wavelength of 620 nm. Each samples were performed in triplicate and average obtained.

### Statistics

Mann-Whitney test (unpaired non-parametric data) and Wilcoxon matched-pairs signed rank test (paired non-parametric data) was used to compare two groups. One-way ANOVA and Tukey’s multiple comparison test (parametric data) or Kruskal-Wallis test with Dunn’s multiple comparison test (non-parametric data) were used for comparison of data with more than two groups. Two-way ANOVA was used to analyze differences between two or more groups over time. Dunnett’s multiple comparison test (to compare with control group) or Bonferroni multiple comparison test (to compare all groups) were used to assess differences at each time points. A P value of <0.05 was taken to be statistically significant. Analysis was carried out using GraphPad Prism 8.2.0 (272).

## Notes

### Competing Interest Statement

The authors have declared no competing interest.

