## Supplemental Figure 1 for "NRP1-expressing monocyte/macrophages promote arteriogenesis in the ischemic limb"

### Supplementary figures and legends

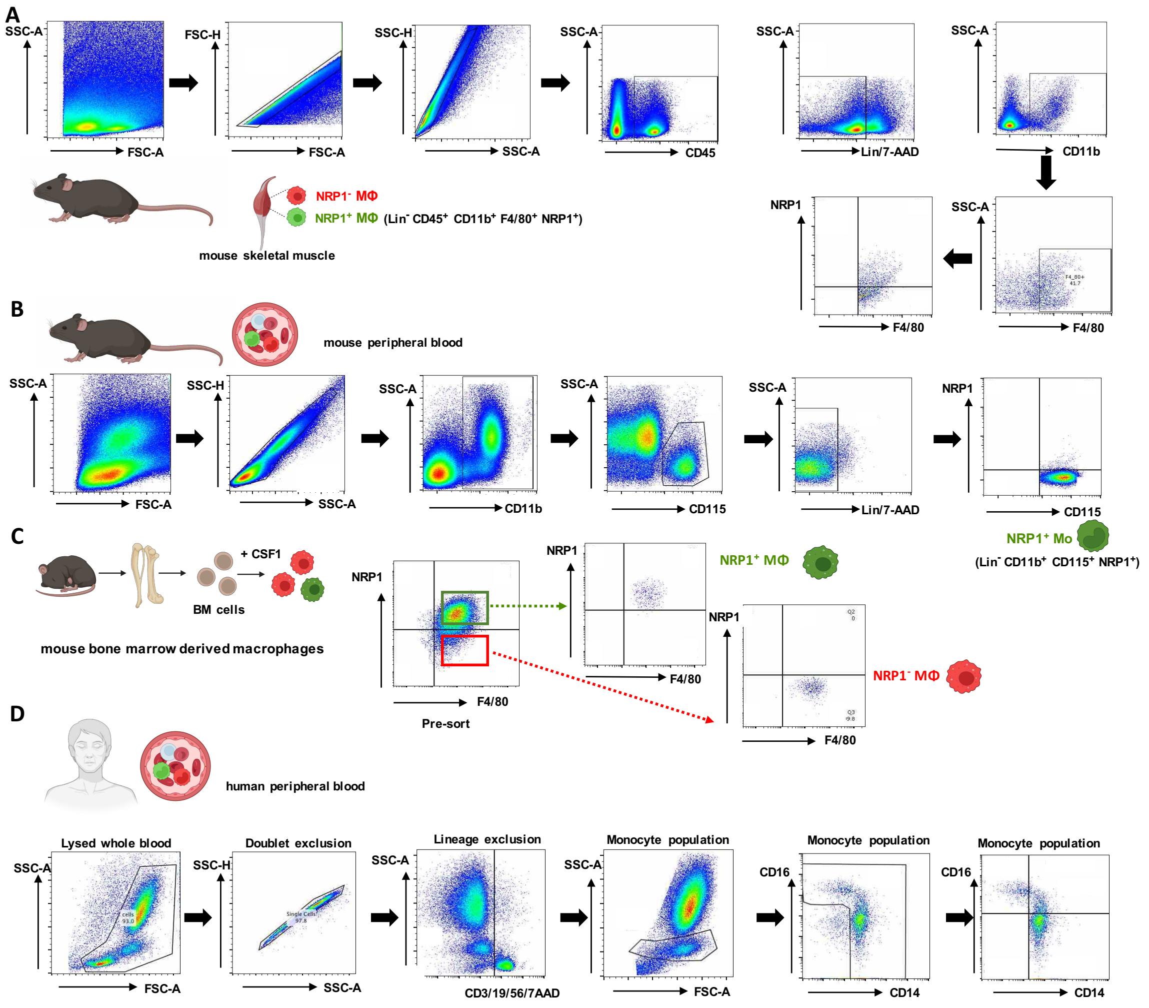


**Fig. S1. Flow cytometric gating strategies for cell analysis and cell sorting.**

**(A)** Flow cytometric gating strategy of tissue macrophages from mouse skeletal muscle.

**(B)** Flow cytometric gating strategy of circulating monocytes from mouse blood.

**(C)** Flow cytometric dot-plot showing the FACS strategy to isolate NRP1$^{+}$ and NRP1$^{-}$ BMDMs.

**(D)** Flow cytometric gating strategy of circulating monocytes in CLI patients.
